# Cerebral perfusion imaging: Hypoxia-induced deoxyhemoglobin or gadolinium?

**DOI:** 10.1101/2021.11.08.467772

**Authors:** E.S Sayin, J. Schulman, J. Poublanc, H. Levine, L. Venkatraghavan, K. Uludag, J. Duffin, J.A. Fisher, D.J. Mikulis, O. Sobczyk

## Abstract

Assessment of resting cerebrovascular perfusion measures (mean transit time, cerebral blood flow and cerebral blood volume) with magnetic resonance imaging currently requires the intravascular injection of the dynamic susceptibility contrast agent gadolinium. An initial comparison between hypoxia-induced deoxyhemoglobin and gadolinium was made for these measures in six healthy participants. A bolus of deoxyhemoglobin is generated in the lung via transient hypoxia induced by an available computer-controlled gas blender technology employing sequential gas delivery (RespirAct™). We hypothesised and confirmed perfusion measures from both susceptibility contrast agents would yield similar spatial patterns of cerebrovascular perfusion measures. We conclude that hypoxia-induced deoxyhemoglobin, an endogenously, non-invasively generated, non-allergenic, non-toxic, recyclable, environmentally innocuous molecule, may be suitable to become the first new magnetic resonance imaging susceptibility contrast agent introduction since gadolinium.

## Introduction

Many common conditions such as cigarette smoking, high blood cholesterol, obesity, sedentary lifestyle, diabetes, hypertension, and aging result in silently accumulating cerebrovascular pathologies, for example small vessel disease, venous collagenosis, chronic inflammation and multiple subcortical infarcts. The accumulation of cerebral vascular disease leads to cognitive decline and can precipitate sudden strokes^1^. The health of cerebral perfusion can be assessed by magnetic resonance imaging (MRI)-based perfusion measures such as mean transit time (MTT), cerebral blood volume (CBV) and cerebral blood flow (CBF). In contrast to flow measures, surrogate measures such as those obtained from cognitive tests^1^ may be affected by vascular function, but are not a direct reflection of it.

MRI-based measures of cerebral vascular perfusion measure require an intravascular injection of a dynamic susceptibility contrast agent. Currently gadolinium (Gd) is the only option, but is invasive, expensive^2^, leaks into the extracellular fluid^3^, accumulates in tissue^4^, has risks of causing harm^5–7^, and is an emerging environmental pollutant^8^. Consequently, there remains an urgent world-wide need for an MRI contrast agent that is non-invasive, economical, remains in the blood pool, does not accumulate in the body, is abundant, safe, and is environmentally innocuous. These challenging requirements may be addressed by hypoxia-induced deoxyhemoglobin (dOHb)^9,10^.

Several a-priori considerations prompted the exploration of dOHb as a potentially useful susceptibility contrast agent. *First*, although oxyhemoglobin is diamagnetic, it becomes paramagnetic after giving up its oxygen to become dOHb^11^. Thus, like Gd, dOHb changes the homogeneity of a magnetic field inside and outside the blood vessels in proportion to its concentration ([dOHb])^12^. *Second*, dOHb can be rapidly formed in the blood of pulmonary capillaries by breathing hypoxic gas as previously shown by Vu et al^13^ and this process can be rapidly reversed by restoring normoxia by a single breath containing normal levels of oxygen^10^. *Third*, the limiting factors in the rate of formation and clearing of dOHb are those in changing the end tidal partial pressure of oxygen (P_ET_O_2_) in the lung. We have developed considerable expertise in inducing rapid, controlled changes in the P_ET_O_2_ using techniques described as *prospective end tidal targeting*^14,15^ and *sequential gas delivery* (SGD)^16,17^. *Finally*, at 3 Tesla, ~25% dOHb in arterial blood provides a T2* signal change sufficient to generate calculations of MTT, CBV and CBF whose magnitudes and distributions are consistent with those previously published for Gd-based perfusion imaging^10^.

Here we describe the use of hypoxia-induced dOHb as a susceptibility contrast agent. We use the ensuing changes in cerebral blood oxygen level dependent (BOLD) signal to calculate cerebrovascular MTT, CBV and CBF. To initially evaluate whether dOHb would be a suitable contrast agent for computing perfusion measures, we compare these measures to those calculated from an intravenous injection of Gd in the same 6 healthy participants. We hypothesize that the relative magnitude and distribution of the perfusion measures calculated using dOHb as a contrast agent will be similar to those generated using Gd.

## Methods

### Participant and Ethics Approval

This study conformed to the standards set by the latest revision of the Declaration of Helsinki and was approved by the Research Ethics Board of University Health Network according to the guidelines of Health Canada. All participant data was anonymized according to institutional protocols. We recruited 6 participants (5 M) between the ages of 23 and 60 by advertisement and word of mouth. All participants provided written and informed consent prior to scanning. All were healthy individuals, non-smokers, not taking any medications, and with no history of neurological, cardiovascular or kidney diseases.

### Respiratory Protocol

A computerized sequential gas delivery system, (RespirAct™, Thornhill Medical, Toronto, Canada) was used to control and manipulate P_ET_O_2_ while maintaining normocapnia, independent of participant ventilation^18,19^. Participants breathed through a facemask sealed to the face with skin tape (Tegaderm, 3M, Saint Paul, MN, U.S.A.) to exclude all but system-supplied gas. The programmed P_ET_O_2_ stimulus pattern was 4-minutes and 20 seconds long (Fig. 1), and consisted of a 60 s baseline P_ET_O_2_ of 95 mmHg (normoxia), a step decrease in P_ET_O_2_ to 40 mmHg (hypoxia) for 60 seconds, a return to normoxia for 20 seconds, a step decrease in P_ET_O_2_ to 40 mmHg for 60 seconds, followed by a return to normoxia for 60 s. Arterial [dOHb] was calculated from P_ET_O_2_ using the Hill equation describing the *in-vivo* oxyhemoglobin dissociation curve^20^.

**Figure 1.**
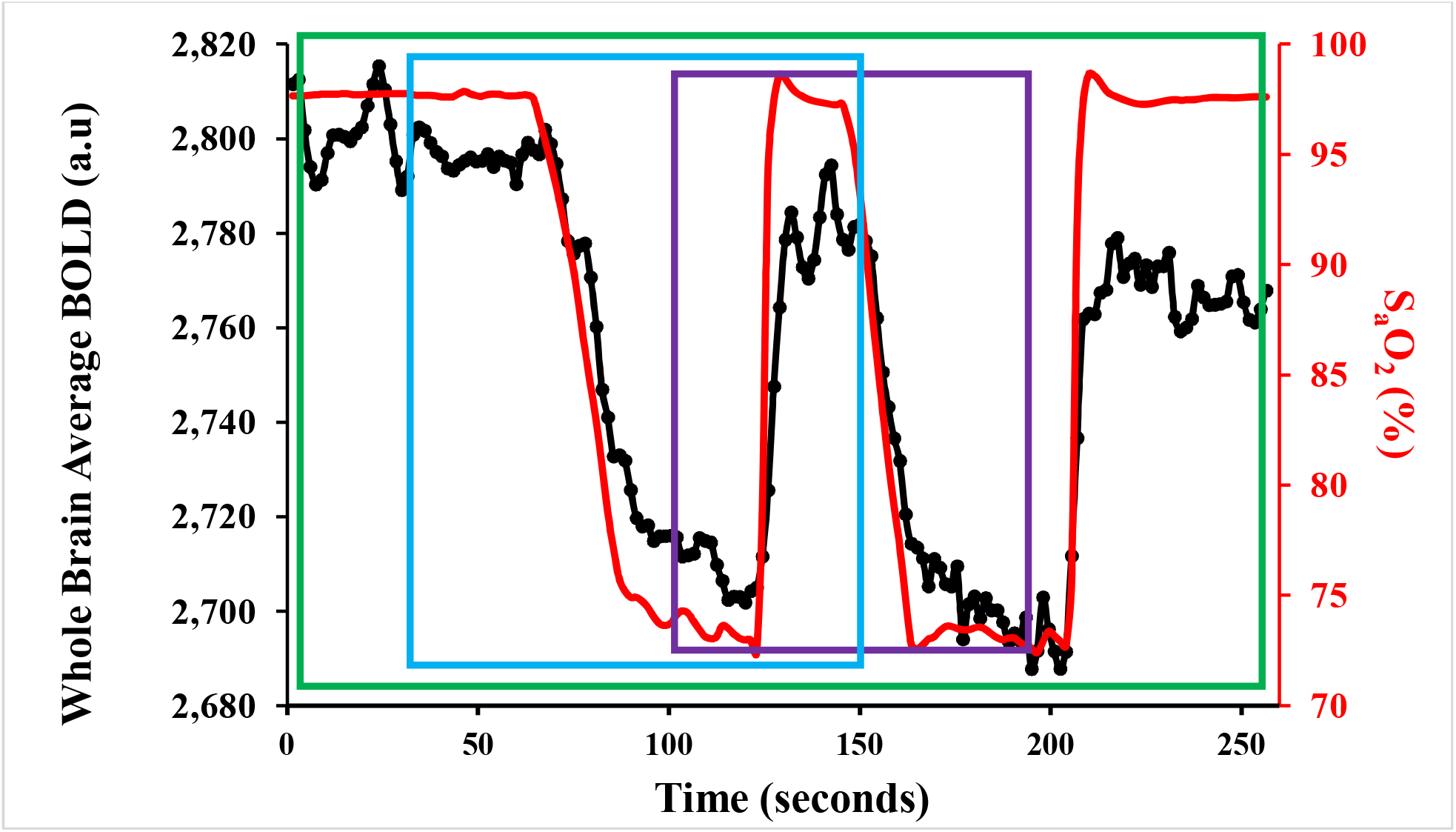
The test protocol in a representative participant. The arterial O_2_ saturation (S_a_O_2_) (red), calculated from the P_ET_O_2_ and the oxygen dissociation curve^20^, and the whole brain average BOLD response (black) in arbitrary units (a.u). The test protocol was divided into selected parts for separate analyses: “single hypoxic” (blue rectangle), “double hypoxic” (green rectangle) and “single normoxic” (purple rectangle).

### MRI Scanning Protocol

The study was performed using a 3-Tesla scanner (HDx Signa platform, GE healthcare, Milwaukee, WI, USA) with an 8-channel head coil. The experimental protocol consisted of a high-resolution T1-weighted scan followed by two BOLD sequence scans on all participants. The first BOLD scan sequence was acquired during P_ET_O_2_ manipulation, while the second one was acquired during an injection of Gd. The high-resolution T1-weighted scan was acquired using a 3D spoiled gradient echo sequence with the following parameters: TI = 450 ms, TR 7.88 ms, TE = 3ms, flip angle = 12°, voxel size = 0.859 × 0.859 × 1 mm, matrix size = 256 × 256, 146 slices, field of view = 24 × 24 cm, no interslice gap. The BOLD data was acquired using a T2*-weighted gradient echoplanar imaging sequence with the following parameters: TR = 1500 ms, TE = 30 ms, flip angle = 73°, 29 slices voxel size = 3 mm isotropic voxels and matrix size = 64 × 64. After the completion of P_ET_O_2_ targeting protocol, the participant returned to free breathing on room air for at least 5 minutes before the Gd based perfusion acquisition for which Gadovist, 5 ml was injected intravenously at a rate of 5 ml/s followed by 30 ml of saline at a rate of 5 ml/s.

### Data Analysis

#### Basics

MR images generated by changes in P_ET_O_2_ data were analyzed using Analysis of Functional Neuroimaging (AFNI) software (National Institutes of Health, Bethesda, Maryland)^21^. BOLD images were volume registered, slice-time corrected and co-registered to the anatomical images. The T1-weighted spoiled gradient images were segmented into gray (GM) and white matter (WM) using Statistical Parametric mapping 8 (SPM8) (Wellcome Department of Imaging Neuroscience, Institute of Neurology, University College, London, UK).

#### Determining the BOLD signal change and the Contrast to Noise Ratio

The BOLD signal change (ΔBOLD) and contrast to noise ratio (CNR) was calculated for Gd and each selected part of the protocol. To examine the effectiveness of using different parts of the dOHb protocol, calculations were repeated using each selected part of the protocol outlined in Fig. 1.

To calculate CNR, a 20s continuous baseline signal at the beginning or end of each dataset was selected. A linear regression between each voxel BOLD signal and the average whole brain BOLD signal was calculated. The slope of the regression multiplied by the average whole brain BOLD signal peak-to-peak was used to calculate relative ΔBOLD for each voxel of the brain. CNR was calculated by dividing ΔBOLD by the standard deviation of the linear regression residuals^10^. Average CNR and ΔBOLD values for GM, WM and GM/WM ratios were calculated for each participant.

#### Determining the arterial input function for dOHb

Perfusion measures for dOHb were calculated using two separate global arterial input functions (AIF): the measured BOLD signal over the middle cerebral artery (AIF-MCA) and the SaO_2_ (AIF-SaO_2_) calculated from the P_ET_O_2_. Identifying the AIF from a voxel over the MCA is fraught^22^; however, we expended our best efforts to optimize the choice of voxel. Smoothing was applied to each voxel in the dataset using an adaptive mean filter of width (n=7) using AFNI software. FSL tool VERBENA software ^23^ was used to calculate MTT and relative CBF (rCBF) using the Gd data and the data from the double hypoxic protocol using each AIF. Relative CBV (rCBV) was calculated as MTT multiplied by rCBF. Maps of the dOHb and Gd perfusion measures were generated using AFNI software and overlayed onto their respective anatomical images. To normalize for rCBF and rCBV perfusion maps within participants for Gd, dOHb (MCA), dOHb (SaO_2_), the ratio between whole brain average Gd and the respective whole brain average for each dOHb AIF were calculated. Furthermore, the dOHb perfusion maps were scaled with the calculated ratio to normalize the relative measures within participants. Average MTT values (the only absolute measure) for GM, WM, and GM/WM ratios were calculated for each participant.

#### Statistical Analysis

Statistical comparisons within participants for CNR and ΔBOLD values across all methods (Gd and each protocol) for GM, WM and GM/WM ratios were performed using a one-way repeated measures analysis of variance (rmANOVA) with a Holm-Sidak method of multiple comparisons correction (*α* = 0.05). In the event of an Equal Variance test or a Normality Test (Shapiro-Wilk) failure, a Friedman Repeated Measures Analysis of Variance on Ranks was used with Tukey Test method of multiple comparisons correction. A rmANOVA was performed on the average MTT values for GM, WM and GM/WM ratios between Gd (AIF-MCA), dOHb (AIF-MCA) and dOHb (AIF-SaO_2_) for each participant (*α* = 0.05).

## Results

### The Hypoxic Stimulus

The brief (60 s) hypoxia induced in these tests was well tolerated by all participants. After completion of the test, only one of the participants reported having had moderate shortness of breath.

### The BOLD Signal

The ΔBOLD and CNR for WM and GM are presented in Table 1 for Gd and the three selected parts of the protocol: single hypoxic, double hypoxic, and single normoxic. Fig. 2 displays axial images of ΔBOLD maps in a representative participant. Although all types of the selected parts of the protocol yielded similar values, the double hypoxic part resulted in the highest ΔBOLD and CNR (Table 1). Consequently, we chose the double hypoxic part in the calculations of the perfusion measures for comparison to Gd.

**Figure 2.**
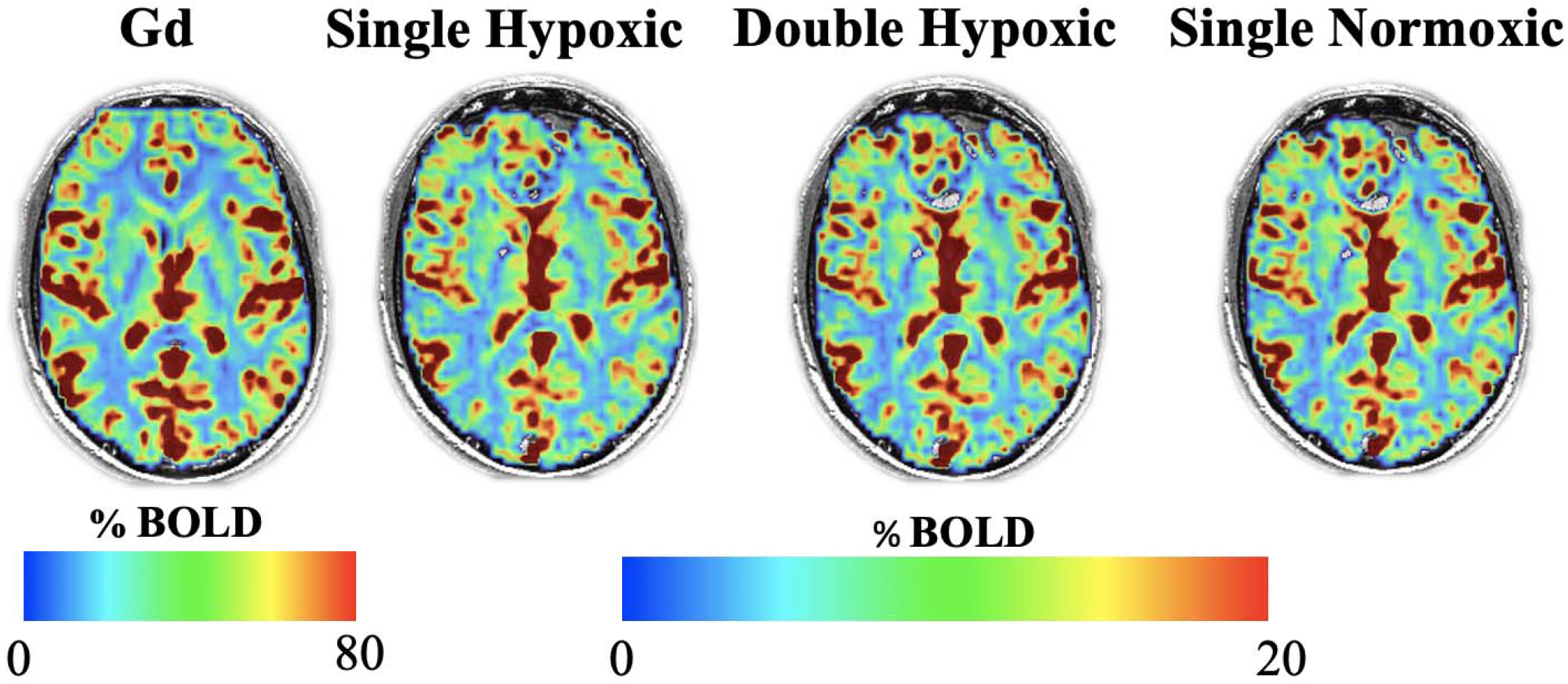
The ABOLD maps for a representative participant comparing Gd and selected parts of the protocol: single hypoxic, double hypoxic, and single normoxic. Note: If the difference in signal strength is accounted for, all the maps appear similar.

**Table 1.**
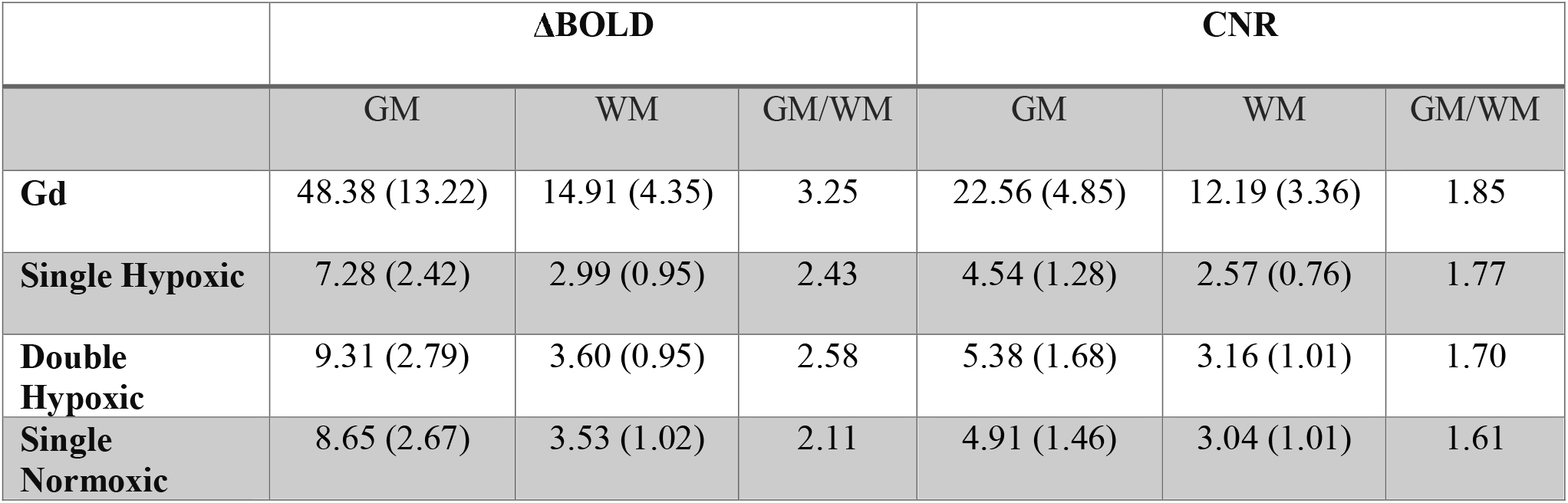
Average (SD) measures of ΔBOLD and CNR calculated for Gd, and the three selected parts of the protocol: single hypoxic, double hypoxic, and single normoxic for the six participants.

| | ΔBOLD | | | CNR | | |
| --- | --- | --- | --- | --- | --- | --- |
|  | GM | WM | GM/WM | GM | WM | GM/WM |
| **Gd** | 48.38 (13.22) | 14.91 (4.35) | 3.25 | 22.56 (4.85) | 12.19 (3.36) | 1.85 |
| **Single Hypoxic** | 7.28 (2.42) | 2.99 (0.95) | 2.43 | 4.54 (1.28) | 2.57 (0.76) | 1.77 |
| **Double Hypoxic** | 9.31 (2.79) | 3.60 (0.95) | 2.58 | 5.38 (1.68) | 3.16 (1.01) | 1.70 |
| **Single Normoxic** | 8.65 (2.67) | 3.53 (1.02) | 2.11 | 4.91 (1.46) | 3.04 (1.01) | 1.61 |

*GM Gd CNR was significantly different from those of all parts of the protocol (P<0.001), however no significant differences found among the parts (P>0.8). WM Gd CNR was significantly different from those of the single hypoxic and normoxic parts (P<0.05) but not the double hypoxic part (P>0.05). The only significant difference in CNR was found in the GM/WM ratio between Gd and that of the single normoxic part (P<0.05). The GM ΔBOLD Gd was significantly different from those of all parts (P<0.001) which did not differ among themselves (P>0.8). In addition, WM ΔBOLD Gd was significantly different from those of all parts (P<0.001) however no significant difference was found between parts (P>0.8). The only significant difference in ΔBOLD GM/WM ratio was between Gd and the single hypoxic and normoxic parts (P<0.05).*

### Determining the AIF

The perfusion measures, MTT, rCBF and rCBV, were calculated for Gd using an AIF from the MCA. Similarly, they were calculated for the dOHb double hypoxic part using an AIF from both the MCA and SaO_2_. Fig. 3 displays perfusion maps for a representative participant. The perfusion maps for all participants can be found in the supplementary data (Fig. S1).

**Figure 3.**
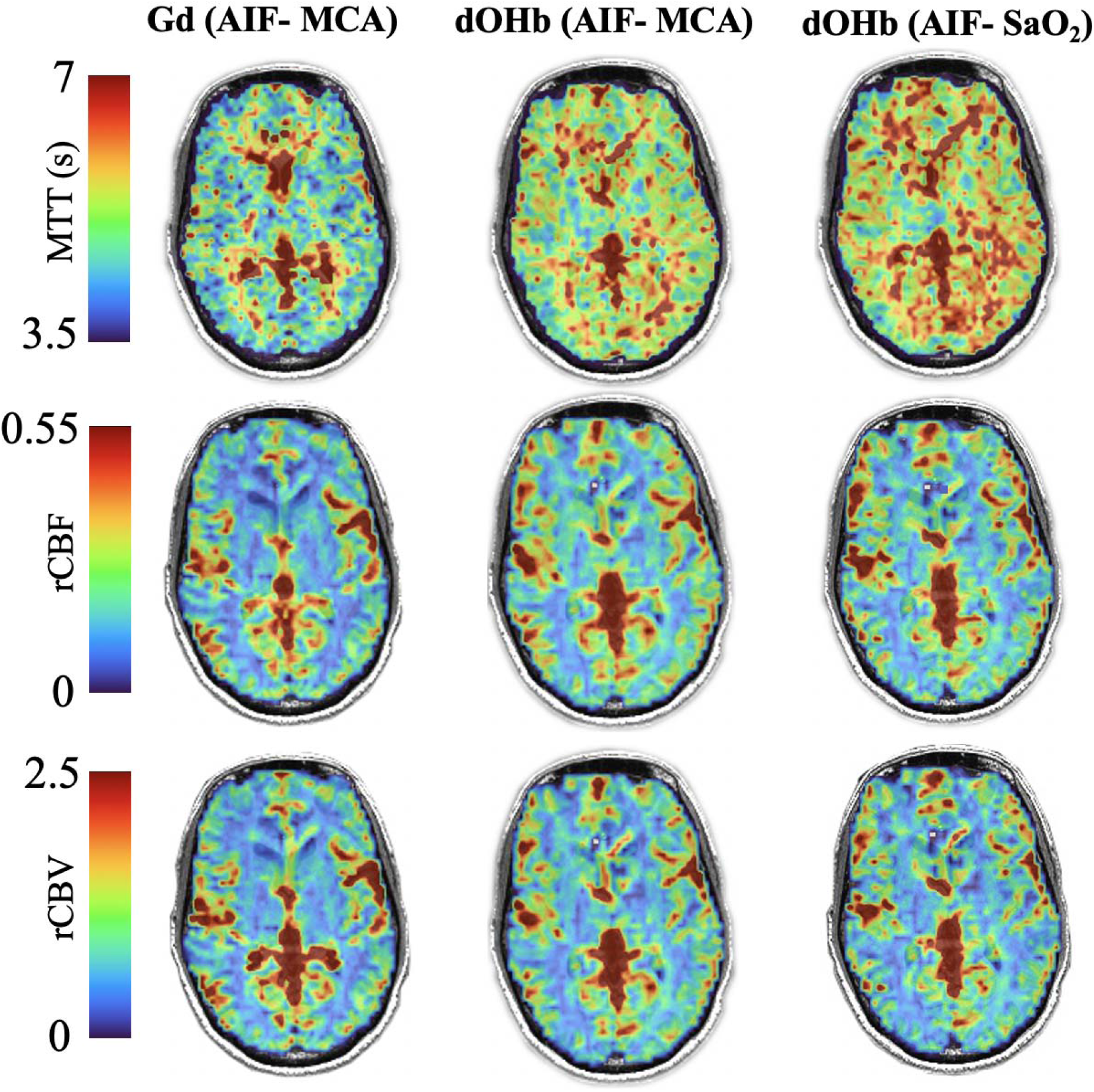
Perfusion maps of MTT, rCBF and rCBV of a representative participant comparing dOHB: AIF-MCA and AIF-S_a_O_2_ to Gd. Note: The spatial distribution of dOHb AIF-MCA and AIF-SaO_2_ are similar to each other and Gd AIF-MCA.

### MTT

The average MTT, the only measure in absolute units, in GM and WM for Gd and the double hypoxic part of the protocol using both the MCA and SaO_2_ AIF in 6 participants is displayed in Fig. 4. There were no significant within-participant differences in MTT GM/WM ratio (p=0.282, α =0.05).

**Figure 4.**
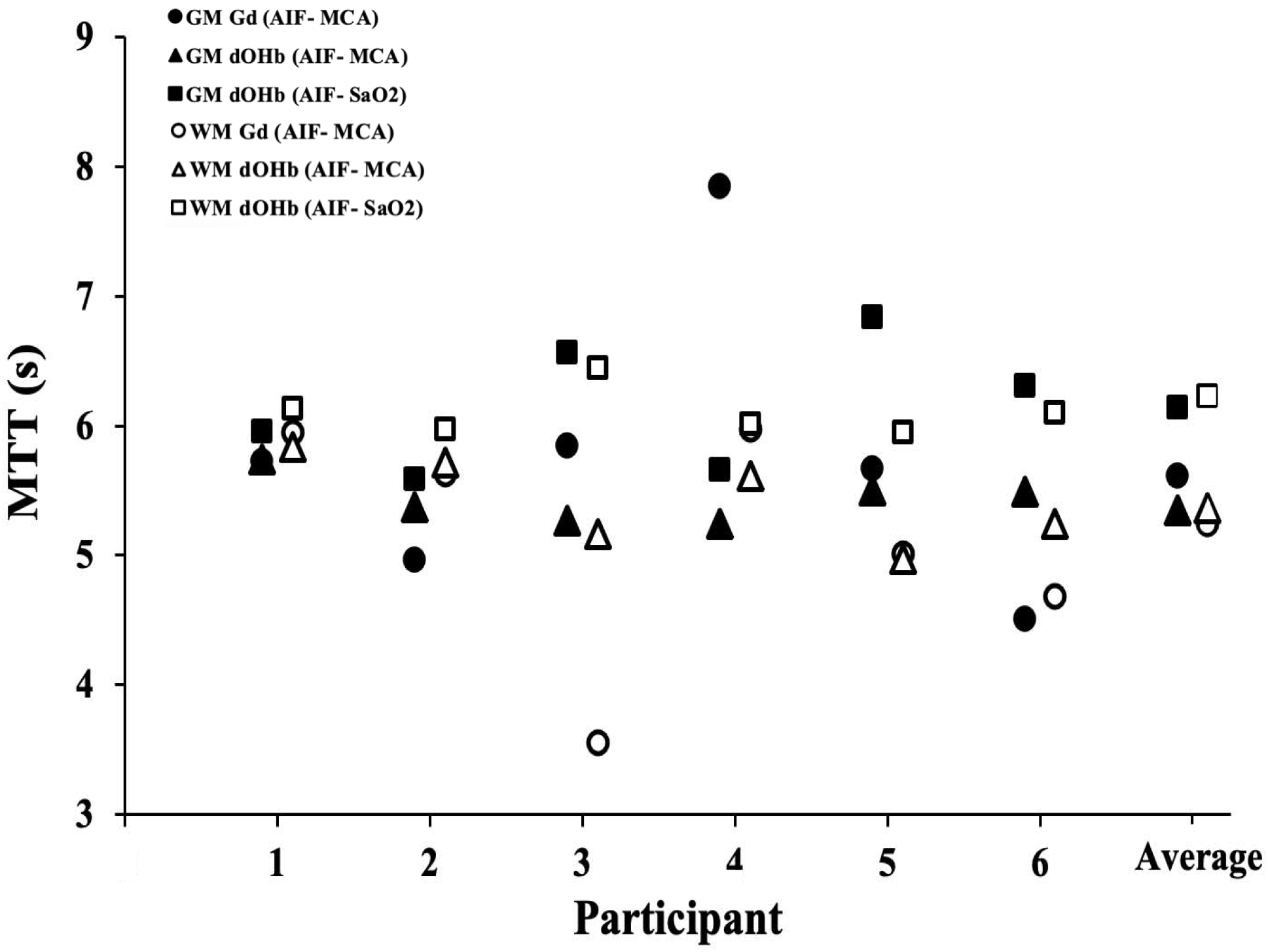
MTT values (s) for Gd and dOHb in GM and WM for the 6 participants. The average values are shown on the right. A rmANOVA found that MTT measures within each participant were not significantly different in GM (P = 0.325). The following comparisons were not significantly different: WM, MTT measures calculated for dOHb (AIF-MCA) and (AIF-SaO2) (P = 0.177); MTT measures for Gd (AIF-MCA) and dOHb (AIF-MCA) (P = 0.384). However, within participant MTT differences between Gd (AIF-MCA) and dOHb (AIF SaO_2_) were small but statistically significant (P = 0.038).

## Discussion

### Main finding

The main finding of this preliminary study is that dOHb used as a contrast agent displayed similar pattern and spatial distribution in maps of MTT, rCBV and rCBF to those generated by Gd. Currently, Gd is the only option for consistent susceptibility-based contrast imaging. By comparing perfusion measures obtained using Gd and dOHb in the same participants under uniform protocols we sought to control for methodological, temporal, and individual participant variables, and thereby isolate differences in results as indicators of inherent differences between the two methods. Despite minimal effort at optimization of dOHb-based data, these differences were judged to be small and justify follow-up investigation. The overarching aim would be to recruit dOHb contrast as a non-invasive, repeatable, rapid, direct measure of resting cerebrovascular perfusion suitable for monitoring the status and progress of cerebrovascular pathophysiology underlying condition such as cognitive dysfunction and stroke.

### Hypoxia tolerance and safety

In designing this protocol, we estimated that a physiologically tolerable arterial PO_2_ of about 40 mmHg would produce a [dOHb] change of about 25%^20^ which would generate a sufficient change in the BOLD signal at 3T to test the value of dOHb as a susceptibility contrast agent^10^. To put the hypoxia used in these tests into perspective, a study of the cerebral blood flow effects of hypoxia reported that test participants had no distress and negligible changes in respiratory rate and heart rate while PO_2_ was maintained at about 50 mmHg for up to 20 min^24^. Rebreathing tests at isoxic P_ET_O_2_ = 40 and 50 mmHg are routinely employed for assessing respiratory chemoreflexes^25^ even in heart failure patients ^26^. Finally, we note that some 20-40 million Americans suffer from moderate to severe obstructive sleep apnea, many of whom also have severe comorbidities including cerebral and myocardial ischemia^27^, and sleep studies have recorded the frequent occurrence of obstructions where P_ET_O_2_ falls into the 20-40 mmHg range^28^.

### Methodological Considerations

This study followed the recently described findings of Coloigner^29^ and Vu^13^ who demonstrated that abrupt increases in arterial [dOHb], induced by transient hypoxia, produce measurable changes in the BOLD signal. Their reports noted the similarity of perfusion measures using dOHb as a contrast agent to historical values but did not validate the relationship by comparison to measures using Gd. Abrupt transient changes in [dOHb] have not otherwise been considered as a susceptibility contrast agent due to the complicated nature of precisely controlling and inducing abrupt targeted changes in P_ET_O_2_ in individuals. Vu and colleagues used inhaled nitrogen, a popular technique to produce a transient hypoxia ^30^. However, when using nitrogen as the source gas, and a 2 L reservoir on the inspiratory limb to buffer the progress of hypoxia, the duration and magnitude of the reduction of P_ET_O_2_ become highly sensitive to the pattern and extent of breathing, and therefore lacked precise control^14^.

In this study, we examined the CNR for the three selected parts of the protocol. Previously, Poublanc et *a*l^10^ determined that CNR increased with repeated hypoxic stimuli but minimal improvement after a second hypoxic stimulus. The results from our experiments showed that the double hypoxic part of the protocol yielded a slightly larger CNR compared to the single hypoxic or the single normoxic parts (Table 1), consequently that was the test protocol used for comparison to Gd. The differences between the three protocols were small indicating that further studies may yield a suitable compromise between a simpler protocol and CNR.

For Gd, abrupt changes in concentration are attempted by rapid intravenous injection at 5 ml /s followed by a bolus of saline. Nevertheless, the contrast is dispersed as it passes the right heart, loses contrast in the lungs and mixes with the blood in the left heart. To approximate a squarewave leading edge for a dOHb bolus it is necessary to implement a rapid change of P_ET_O_2_ in the lung, which is resisted by the dispersing effects of the inspired gases progressively diluting and replacing those in the lungs of participants. The approach used here to induce a targeted level of hypoxia was to pre-calculate and administer the optimum level of nitrogen and oxygen for each successive breath to most efficiently attain and maintain a targeted P_ET_O_2_, taking into account the lung volume, oxygen consumption, and breath size. We emphasize that the great advantage of these methods is that the calculations are made prospectively and do not rely on feedback loops, so that the trajectory to the target P_ET_O_2_ is optimized with each breath^14^. The delivery of the optimized gases is automatically synchronized to the particular breathing pattern of the participant by an automated gas blender (RespirAct™) implementing sequential gas delivery^17^. The rapidly formed dOHb in the pulmonary capillaries traverses only the left atrium and left ventricle, minimizing the opportunity for dispersion in the blood before passing into the cerebral arteries. We suggest this accounts the high similarity of the perfusion measures calculated from AIF’s using P_ET_O_2_ and voxel BOLD signal over the MCA.

### Differences between Gd and dOHb

As an intravascular paramagnetic molecule, dOHb has several characteristics that differ from those of Gd (Supplementary Data Table 1). *First*, while Gd distributes throughout the plasma, dOHb is confined inside red blood cells. One consequence of this differing distribution is that [Gd] increases as the hematocrit falls in the capillaries while [dOHb] increases^31^. These effects have implications as to the relationship between signal size and contrast concentration^32^. A second consequence is that Gd passes readily out of capillaries into tissues such as the lungs resulting in dispersal before arrival in the arteries. In the brain, Gd is contained intravascularly by the blood-brain barrier making it a good marker for its integrity, for example, in brain tumors^33^. By contrast, dOHb does not leak into tissues and remains intravascular regardless of the blood-brain barrier unless there is active bleeding. *Second*, the arterial [Gd] varies with dose^34^, speed of injection^35^, cardiac output^36^, degree of dispersion^37^, and amount leaked from the vessels prior to arrival at the arteries^38^. Of these, only dose and speed of injection can be controlled, so that it is not possible to deliberately implement a particular concentration or profile of [Gd] as an AIF. An AIF may be measured at the MCA, but this measurement is also problematic^39,40^, and in most cases must be deconvolved from the residue function. Whereas the exact [dOHb] and its profile corresponds to that of P_ET_O_2_. It is independent of cardiac output and existing leaks into interstitial spaces. *Finally*, for Gd, calculations of the cerebral perfusion measures must be measured on the first pass of Gd as some of the molecules traversing high flow short distance organs such as thyroid, heart and bronchial circulation may recirculate quickly^35^, with its arrival potentially overlapping the AIF. Gd is only slowly cleared from circulation, thereby confounding further measures for about 2 hours^41^. By contrast, all changes in dOHb are oxidized when P_ET_O_2_ is restored to normoxia, enabling multiple repeated measures at close time intervals.

### Limitations

Optimization of the dOHb method is ongoing. We believe that the selected hypoxic stimulus pattern used in this study generated an adequate bolus of contrast as indicated by the agreement of the dOHb perfusion maps with those generated using Gd. However, further development is needed to determine the optimal [dOHb] and the AIF pattern for studies of people with healthy vasculature, as well as those with pathology. The scan sequence parameters need to be optimized for whole brain coverage, temporal resolution (TR) and thereby CNR. In addition, the selection of the AIF in an MCA voxel compared to that using the SaO_2_ requires further investigation.

### Future Directions

For this study we chose a particular hypoxic stimulus pattern to induce the dOHb changes, and magnetic resonance scanning settings that we felt were appropriate. However, these choices may be optimised after further experimentation. We compared perfusion measures between dOHb and Gd, but the considered gold-standard for such measures uses positron emission tomography (PET). Comparisons of Gd and PET show only moderate consistency^42–44^. Nevertheless, it would be instructive to compare the maps of perfusion measures against this gold-standard, particularly in patients with cerebrovascular disease. This study was an initial comparison of a new susceptibility molecule to a well-known clinical standard. Larger studies are required to identify normal ranges of perfusion measures using dOHb as a contrast agent in population pools divided by age, sex, underlying diseases, and specific cerebrovascular pathophysiology.

## Conclusion

In conclusion, using dOHb as a contrast agent provided similar set of perfusion maps and measures to those of Gd in the same healthy individuals. This finding supports the continued exploration of dOHb as a finely controlled endogenous contrast agent with the advantages of being an endogenous, non-invasively generated, non-allergenic, non-toxic, recyclable, environmentally innocuous molecule. It may be the first new MRI susceptibility contrast agent introduction since gadolinium.

## Data Availability

Anonymized data will be shared by request from any qualified investigator for purposes such as replicating procedures and results presented in the article provided that data transfer is in agreement with the University Health Network and Health Canada legislation on the general data protection regulation.

## Acknowledgements

All the authors thank Toronto Western Hospital MR technologist Keith Ta for all his help in acquiring the MR imaging data, Abby Skandaraniyam for study coordination and Dr. John Wood for feedback on the final draft of the paper. The study was supported by the Institute for Basic Science, Suwon, Republic of Korea (IBS-R015-D1) to Kamil Uludag.

## Author contributions

All authors contributed to the design and conceptualization of the study. ESS, OS, JD and JP were involved in the data analysis. ESS, OS, JP, drafted the initial draft of the manuscript. All authors participated in multiple rounds of review of data and editing the manuscript. All authors have reviewed and endorse the final draft.

---

**Supplementary Figure 1:**
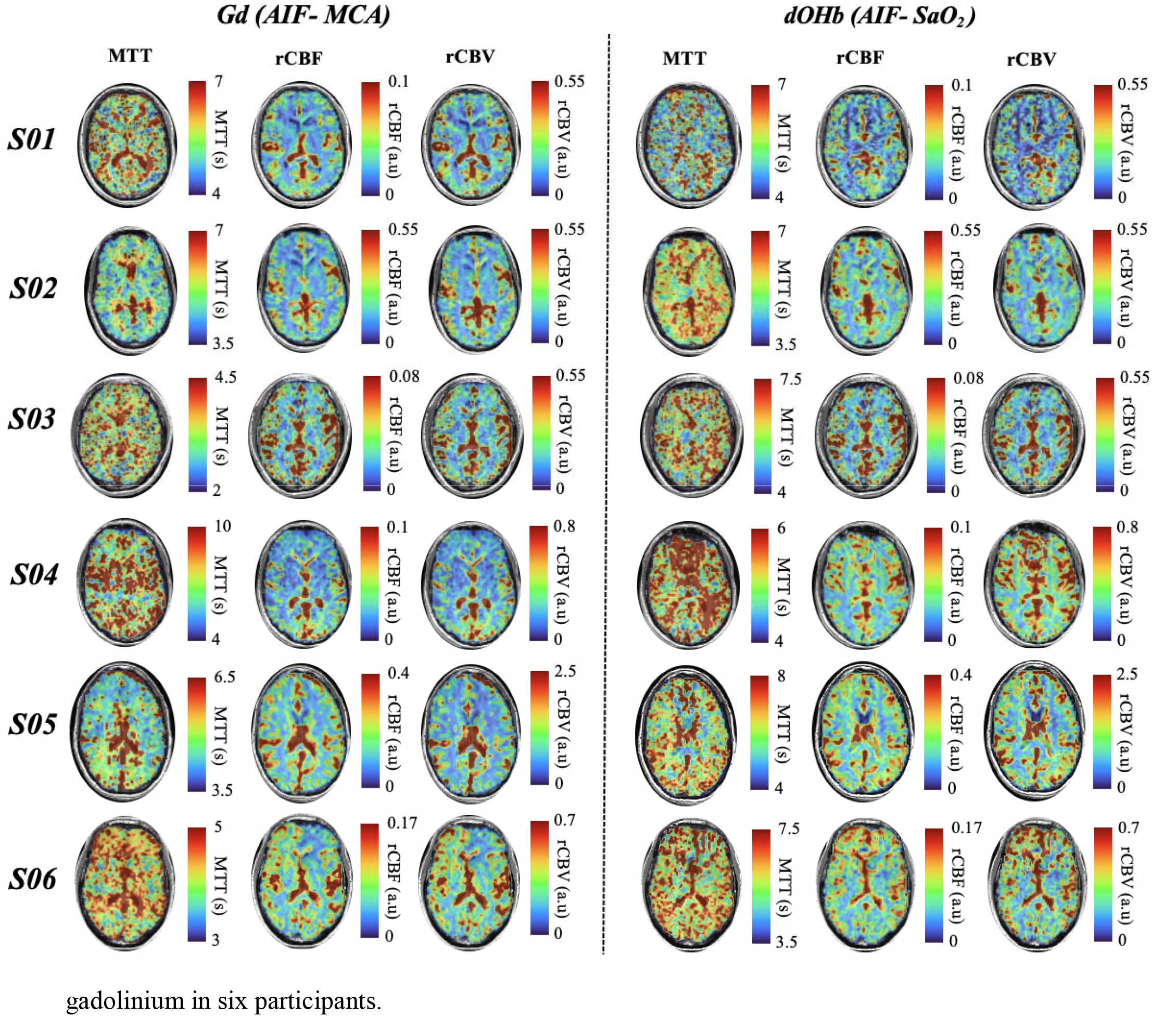
The perfusion maps using dOHb and Gd in all 6 subjects. The perfusion maps showcasing similarities of the two different methods, deoxyhemoglobin, and gadolinium in six participants.

**Supplementary Table 1:** Comparison of Gd based contrast agent and [dOHb] as susceptibility contrast agents. Abbreviations: ([dOHb]) deoxyhemoglobin concentration; ([Gd]) gadolinium concentration; (AIF) arterial input function.

| **Performance characteristic** | **[Gd]** | **[dOHb]** |
| --- | --- | --- |
| **Non-invasive** | NO<br>Intravascular injection | YES<br>Contrast bolus generated by inhaling gas |
| **High SNR: # unpaired electrons** | 7 unpaired electrons<br>The paramagnetic effect is several-folds higher inducing a greater $\Delta T2^*$ | 4 unpaired electrons |
| **Agent tied to physiological environment** | NO<br>(inert) | YES<br>Signal strength proportional to [dOHb] |
| **Contrast remains intravascular** | NO | YES |
| **Contrast concentration known** | NO<br>**Enabling blood [Gd] calculation:**<br>Known dose<br>Known rate of injection<br>**Obscuring blood [Gd] calculation:**<br>Saturation of signal loss possible with excessive dosing.<br>Unknown (i) cardiac output; (ii) dispersion in venous system; (iii) extravasation during lung and heart transit; (iv) dispersion in heart | YES<br>Precise arterial [dOHb] calculated from $P_{ET}O_2$ *<br>Minimal dispersion in healthy left ventricle<br>*Note: Accuracy may be limited by disease with right to left shunt in heart and lungs. |
| **Same scale as tissue curve,** | Relation of arterial and tissue scales uncertain | Intravascular agent only |
| **or known**<br>**scale factor** | • $R2^*$ quadratic dependence on [Gd] |  |
| **signal**<br>**saturation** | • Saturates easily at high doses due to high SNR; phase analysis may also saturate<sup>45</sup> | • Low SNR: Unlikely to saturate |
| **No signal**<br>**aliasing** | • Aliasing if phase shift > 180 deg<sup>45</sup> | • Phase shift analysis unknown |

